# Characterization of a Novel Plasmid in *Serratia marcescens* Harboring *bla*_GES-5_ Isolated from a Nosocomial Outbreak in Japan

**DOI:** 10.1101/2021.07.27.454091

**Authors:** Noriko Nakanishi, Shoko Komatsu, Tomotada Iwamoto, Ryohei Nomoto

**Affiliations:** Department of Infectious Diseases, Kobe Institute of Health, 4-6-5 Minatojima-nakamachi, Chuo-ku, Kobe, Hyogo, 650-0046, Japan

**Keywords:** *Serratia marcescens*, crabapenemase, *bla*_GES-5_, whole-genome sequencing

## Abstract

*Serratia marcescens* is a nosocomial pathogen with carbapenem resistance, limiting the availability of effective treatment options. In this study, we performed molecular characterization of GES-5 carbapenemase-producing *S. marcescens* isolated from an outbreak in Japan. Comparative genetic analysis revealed that the *bla*_GES-5_–encoding plasmid p2020-O-9 is a unique plasmid contributing towards carbapenem resistance. Furthermore, this study highlights the necessity of surveillance programs for monitoring novel, along with commonly occurring carbapenemases in clinical settings.

---

*Serratia marcescens* is a major opportunistic pathogen known to cause nosocomial infections associated with high morbidity and mortality (1, 2). As this bacterium shows resistance towards widely used carbapenem class of antibiotics, treatment options available for nosocomial infections becomes limited (3–5). Carbapenemases (a class of β-lactamases) identified in Japan comprise KPC, GES, NDM, IMP, VIM, and OXA-48 variants, with IMP being the most prevalent in the country (6). GES type Amber class A β-lactamases, in addition to exhibiting β-lactam resistance, also confer carbapenemase activity, such as the GES-5 variant (7, 8). In recent times, nosocomial outbreaks have largely been attributed to GES-5 carbapenemase-producing bacterial strains in various countries including Japan, wherein the first outbreak of *P. aeruginosa* harboring the *bla*_GES-5_ gene occurred in the year 2014 (9–13). This study reports the genetic characteristics of a different GES-5 producing bacterium, *S. marcescens* isolated from an ICU outbreak in Japan in 2020.

A total of six carbapenem-resistant *S. marcescens* strains were isolated from the samples from three ICU patients collected during May-October 2020 (Table 1). Strain was identified using MALDI Biotyper (Bruker Daltonics K.K., Yokohama, Japan), and the minimal inhibitory concentrations (MICs) of various antimicrobial agents were determined by E-test (bioMérieux Japan Ltd., Tokyo, Japan) following the manufacturer’s guidelines. As indicated in Table 1, all of the six *S. marcescens* strains were resistant to imipenem, meropenem, and ceftazidime. Modified carbapenem inactivation method (mCIM) indicated the absence of carbapenemase genes in all the strains. This was supported by the negative results of PCR screening conducted for the major carbapenemase genes *bla*_IMP_, *bla*_NDM_, *bla*_KPC_, and *bla*_OXA-48-like_; however, a positive result was obtained for *bla*_GES_. The MICs of antimicrobial agents (except ampicillin (AM) and cefotaxime (CTX)) in 2020-O-14-2 strain lacking the *bla*_GES-5_ gene, isolated from patient B, were significantly lower than all the other isolates. Whole-genome sequencing of the six isolates harboring *bla*_GES-5_ gene and the one lacking the gene was done for analyzing single nucleotide variations and deriving genetic relationships among the seven isolates to understand the mechanisms of antimicrobial resistance. A survey for analyzing environmental contamination in the ICU was conducted `during the outbreak period`; however, the presence of carbapenem-resistant *S. marcescens* strains was not detected.

**TABLE 1.** Information and the antibiotic susceptibility of the *Serratia marcescens* isolated from three patients involved in the ICU outbreaks

| Patients | Age (yr),<br>sex<sup>a</sup> | No. of isolate | Date of isolation<br>(year/month/day) | Specimens | MICs (μg/ml) for<sup>b</sup> |  |  |  |  |  |  |  |  |  |  |  |  | *bla*<sub>GES-5</sub> | DDBJ accession no.<br>of read data |
| --- | --- | --- | --- | --- | --- | --- | --- | --- | --- | --- | --- | --- | --- | --- | --- | --- | --- | --- | --- |
|  |  |  |  |  | AM | CTX | CAZ | FEP | ATM | IMP | MPM | DOR | ETP | AK | MC | CIP | PMB |  |  |
| A | 76, F | 2020-O-9 | 2020/5/25 | Sputum | >256 | >32 | >32 | 6 | 6 | >32 | >32 | >32 | >32 | >256 | 4 | 0.5 | >1024 | + | DRR308222 |
|  |  | 2020-O-12 | 2020/6/23 | Blood | >256 | >32 | >32 | 3 | 4 | >32 | >32 | >32 | >32 | >256 | 3 | 1 | >1024 | + | DRR308223 |
|  |  | 2020-O-21-2 | 2020/8/3 | Sputum | >256 | >32 | >32 | 2 | 3 | 6 | >32 | >32 | 8 | 64 | 3 | 1.5 | 64 | + | DRR308224 |
|  |  | 2020-O-21-3 | 2020/8/4 | Blood | >256 | >32 | >32 | 2 | 8 | >32 | >32 | 24 | >32 | 96 | 3 | 1.5 | 128 | + | DRR308225 |
| B | 70, M | 2020-O-14-1 | 2020/6/19 | Blood | >256 | >32 | >32 | 4 | 6 | >32 | >32 | >32 | >32 | 128 | 12 | >32 | 96 | + | DRR308226 |
|  |  | 2020-O-14-2 |  |  | >256 | >32 | 2 | 3 | 4 | 1 | 1.5 | 1.5 | 4 | 16 | 3 | 3 | 64 | - | DRR308227 |
| C | 59, M | 2020-O-25 | 2020/10/15 | Ascites | >256 | >32 | >32 | 3 | 8 | >32 | >32 | >32 | >32 | >256 | 3 | 0.38 | >1024 | + | DRR308228 |

<sup>a</sup> F, feamale, M, male.
<sup>b</sup>ampicillin (AM), cefotaxime (CTX), ceftazidime (CAZ), cefepime (FEP), aztreonam (ATM), imipenem (IMP), meropenem (MPM), doripenem (DOR), ertapenem (ETP), amikacin (AK), minocycline (MC), ciprofloxacin (CIP), polymyxin B (PMB)

Genomic libraries of all seven isolates were prepared using the Nextera XT DNA library prep kit (Illumina, San Diego, CA, USA) and sequenced on the iSeq 100 system (Illumina). Additionally, Oxford Nanopore Technologies (ONT) were used for constructing a sequencing library of the strain 2020-O-9 with the help of Rapid Barcoding Kit (SQK-RBK004), and sequenced on a MinION device using flow cell type R9.4.1 (FLO-MIN106D). Hybrid assembly of 2020-O-9 genome was performed using Unicycler v0.4.8 (14), and a circular chromosome and plasmid (p2020-O-9) sequence was obtained. The genome sequences were annotated using DFAST (https://dfast.nig.ac.jp/) and are available in DDBJ (DNA Data Bank of Japan) under the accession numbers AP024847 (chromosome) and AP024848 (plasmid). SNP analysis of the remaining 6 strains was performed taking the genome sequence of 2020-O-9 as a reference, using bwa version 0.7.17, samtools version 1.9, and VarScan v2.4.4 (15–17). The analysis revealed high degree of genetic homogeneity with 0 to 5 SNPs consistent among the seven strains isolated, including GES-5 negative strain 2020-O-14-2. The whole-genome sequencing reads are available in DDBJ Sequence Read Archive under the accession number listed in Table 1. Since MICs of carbapenems in 2020-O-14-2 strain lacking the *bla*_GES-5_ gene were significantly decreased, it is evident that the newly detected *bla*_GES-5_ containing plasmid contributes to carbapenem resistance.

Further genetic characterization of p2020-O-9 was done using ResFinder 3.2 to identify resistance genes, and plasmid incompatibility replicon typing was performed using PlasmidFinder 2.0 developed by the Center for Genomic Epidemiology (http://www.genomicepidemiology.org/). The chromosomal sequence of 2020-O-9 consisted of a single antimicrobial resistance gene *bla*_SRT-2_, as identified by existing databases. p2020-O-9 is a novel 23,921-bp circular untypeable plasmid, with the *bla*_GES-5_ gene located between intI1 of class 1 integron and the *qacEΔ1* and *sul1*, together with a gene of GNAT-family N-acetyltransferase, *aac(6)-29a*. It has a GC-content of 61% and carries 26 protein-coding genes (Fig.1). Sequence analysis using NCBI revealed that the plasmid backbone of p2020-O-9 showed highest similarity with plasmid pCAV1374-16 from *Klebsiella oxytoca* with a query coverage of 59%; however, there was no *bla*_GES-5_ in pCAV1374-16 (Accession no. CP011628). The class 1 integron cassette around the *bla*_GES-5_ gene in p2020-O-9 is similar to that of pMRY16-414SMA_2 from *S. marcescens* (Accession no. LC486677) and the plasmid from *Aeromonas hydrophila* strain WCHAH 01-derived plasmids pGES5 (Accession no. KR014105) (Fig. 1). Plasmid sequences similar to the backbone of p2020-O-9 such as pCAV1374-16, have been registered in databases from a wide range of bacterial species. It is believed that *bla*_GES-5_ was integrated into the class 1 integron cassette of a plasmid like pCAV1374-16 to construct the novel p2020-O-9 plasmid. This suggests that p2020-O-9 could be an untypeable plasmid with the potential to distribute itself widely in *Enterobacteriaceae*.

**Figure 1.**
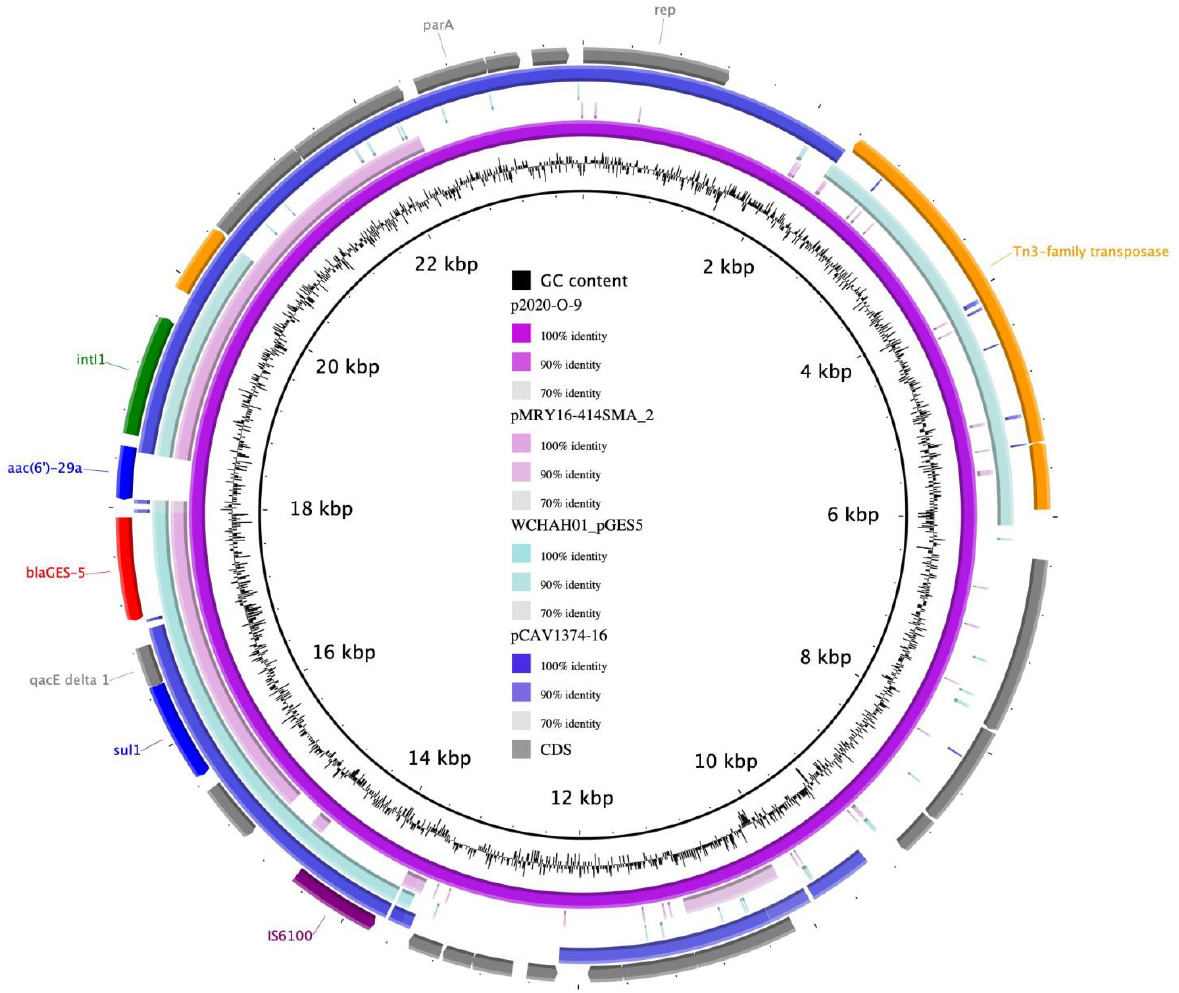
Circular representation of p2020-O-9 was generated using BLAST Ring Image Generator 0.95 (18). A comparison with the pMRY16-414SMA_2 from *Serratia marcescens* MRY-414SMA (LC486677), pGES5 from *Aeromonas hydrophila* WCHAH 01 (KR014105), and pCAV1374-16 from *Klebsiella oxytoca* (CP011628). The outermost circle shows the coding sequence of p2020-O-9. Red, carbapenemase genes; blue, other antimicrobial resistance genes; purple, IS6100; orange, transposase and recombinase genes; green, integrase genes; grey, other genes or coding sequences.

In summary, we reported the molecular characterization of GES-5 producing *S. marcescens* strain harboring a novel *bla*_GES-5_ gene-encoding plasmid, p2020-O-9 isolated from the ICU outbreak in Japan The prevalence and transmission of GES-producing bacteria is largely underestimated due to their rare presence in clinical isolates. However, through this study, we highlight the necessity of surveillance programs for monitoring novel, as well as commonly occurring carbapenemases in clinical settings.

## Abbreviations

WGS: whole-genome sequencing

## Data availability

Sequence data that support the findings of this study have been deposited in DDBJ (https://www.ddbj.nig.ac.jp/) with the accession numbers AP024847 and AP024848.

## Author contributions

NN, RN designed the study methods and wrote the first draft of the manuscript. NN, RN, SK collected the data. NN, RN analyzed the data. TI contributed to the writing of the manuscript.

## Ethical approval

This study was approved by the Ethical Review Committee of the Kobe Institute of Health (approval No. SenR3-9).

## Acknowledgments

We are grateful to Dr. Mari Matsui and Dr. Satowa Suzuki at the National Institute of Infectious Diseases, Japan, for their guidance. This work was partially supported by a grant-in-aid from the Japan Agency for Medical Research and Development (AMED) Research Program on Emerging and Re-emerging Infectious Diseases (grant no. JP21fk0108103).

We declare no conflicts of interest.

